# Aptaswitch/aptakiss adenosine sensor design based on the selection of new DNA/DNA kissing complexes

**DOI:** 10.64898/2026.07.26.740769

**Authors:** Eric Dausse, Nicolas Tourasse, Jean-Jacques Toulmé, Laurent Azéma

## Abstract

Kissing complexes involve interactions between complementary loops of two RNA hairpins. These complexes are continously found in RNA structures and are essential for several biological processes, including the dimerization of retroviral RNA genomes, regulation of plasmid copy number, and small RNA-based control of mRNA translation. We previously identified novel RNA-RNA and RNA-DNA kissing complexes through the in vitro selection of oligonucleotides from synthetic libraries. In order to expand the use of kissing complexes to aptamers and DNA nanotechnology, we describe the selection and characterization of DNA-DNA kissing complexes and their application in the development of DNA adenosine aptasensors. Four DNA/DNA kissing complexes were identified from SELEX and characterized by EMSA and fluorescence anisotropy. The couple dk25/dk26 was further modified to generate an adenosine-triggered aptaswitch, where the addition of adenosine selectively induces the dissociation of loop-loop interactions. This paves the way for a new class of nucleic acid modules, which could be useful for the design of controlled nanostructure assembly and disassembly.

## Introduction

Two complementary loops of RNA hairpins are able to interact through the formation of Watson-Crick base pairs, giving rise to the formation of complexes named kissing complexes^[1]^. This interaction highly depends on the loops’ lengths and sequences, are stabilized by magnesium ions and by additional contribution from the junctions between the stems and the loops^[2]^. Kissing complexes have been identified in natural RNA structures, such as higher order RNA multimers or nanocages^[3]^. Indeed, they regulate numerous biological processes^[4]^ such as plasmid copy number under the control of RNAI/RNAII in E. coli^[5]^, HIV-1 RNA dimerization through the DIS element^[6]^, mRNA translation via the formation of OxyS RNA / FhlA mRNA complex in Escherichia coli^[7]^ or IsoA1 RNA / AapA1 mRNA complex in Helicobacter pylori^[8]^. Many more examples can be found in bacteria and viruses^[9]^. In particular, RNA/RNA kissing interactions are involved in the highly conserved stem loop II of Sars-Cov-2 virus. Kissing interactions have found applications in biotechnology, thanks to the programmable and tuneable nature of RNA. It has been used for the design of artificial scaffolds^[10]^, the controlled assembly of RNA condensates^[11]^, or protonucleus^[12]^.

Initially identified against the TAR hairpin element of HIV-1^[13]^, we have previously obtained RNA/RNA kissing complexes by semi-rationally engineered structured hairpin aptamers likely to interact with stem-loops of the IRES of HCV^[14]^ or by in vitro selection of new motifs from a pool of RNA hairpins containing a randomized apical loops^[15]^.

To expand the development of kissing complexes in DNA nanotechnology, we identified hybrid DNA/RNA kissing complexes and developed structure-switching aptamers^[16]^. Structure-switching aptamers display a conformational change between free and bound states with their cognate target. They are usually obtained by Capture-Selex^[17]^ or from semi-rationally design^[18]^. We have previously developed DNA aptaswitches able to engage a kissing complex with the RNA aptakiss partner, depending on the presence or the absence of the analyte. Interestingly, biosensors using optical^[19]^, electrochemical detection^[20]^ and lateral flow devices^[21]^ were engineered taking advantage of such RNA/RNA^[22]^ or RNA/DNA^[23]^ aptaswitch/aptakiss complexes to detect small molecules. Nevertheless, further development of aptaswitches as sensors would benefit from the selection of DNA-DNA kissing pairs, to improve stability and production cost of the biosensor. Only few DNA kissing complexes were developed for structural studies and nanotechnological applications so far^[24]^.

In this study, we aimed at expanding the repertoire of DNA/DNA kissing complexes and their application as biosensors. We report the in vitro selection of DNA/DNA loop-loop complexes. In a second step, we engineered a DNA aptaswitch as a chimera between the kissing interaction loop and the structure-switching aptamer against adenosine. Such system is the first, to our knowledge, DNA/DNA aptakiss/aptaswitch sensor to be responsive to a small molecule.

## Results and discussion

### Selection of DNA loop-loop complexes by SELEX

In order to identify de novo kissing interactions, we designed the selection of DNA-DNA kissing complexes from a starting single-stranded DNA pool structured in hairpins with randomized apical loop (either 10 or 11 nucleotides) and predefined six base pairs-long stems (Figure 1a); it leads to a potential diversity of about 5.2X106 candidates. The stem was flanked by constant sequences at each end for PCR amplification (two base pairs GA/TC located at the base of the stem were part of the primers; Table S1).

**Figure 1.**
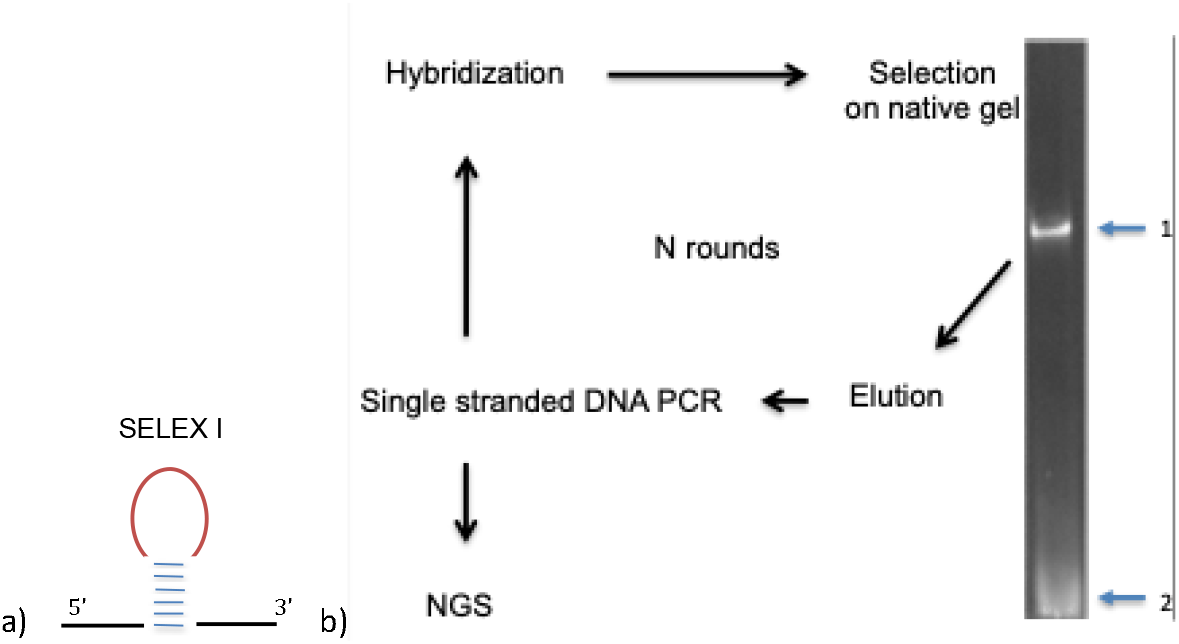
DNA/DNA kissing SELEX design. The library (a) is composed of a random apical loop flanked by constant complementary sequences that form a stem. The stem is extended with constant region required for PCR amplification. (b) SELEX design. The library is first incubated at 25 °C for 12h in order to promote loop-loop interactions. Such complexes are isolated on PAGE native gel : Bound oligonucleotides (arrow 1) were separated from unbound (arrow 2) by Electrophoretic Mobility Shift Assay (EMSA), eluted from the gel and then amplified by PCR. After three rounds, hairpins potentially able to form kissing complexes were deciphered by the Illumina next-generation sequencing (NGS) method.

The SELEX process was designed to promote and isolate loop-loop interactions. The library was heat-denatured and renatured before loading on a non-denaturing polyacrylamide gel. Ethidium bromide-stained gel revealed two bands of interest: the lowest, high mobility band (band 2) corresponding to free candidate hairpins, and an upper band (band 1) with a slower migration which corresponds to two hairpins complexed together, likely engaged in DNA-DNA kissing complexes (Figure 1b). The library (5 µM) was submitted to three selection rounds. Following the first electrophoresis (round 1), performed in a medium containing 10 mM Mg^2+^ to promote kissing interactions^[10d, 25]^, the material from band 1 was extracted and used for the second round after amplification. The second round was actually a negative electrophoretic selection, carried out in the absence of magnesium, to discard DNA duplexes other types of DNA-DNA complexes that might be less impacted by these ionic conditions. Material from band 2, corresponding to Mg-dependent complexes, was extracted, amplified and used (0.5 µM) in round 3. Round 3 was performed as a positive selection with increased stringency (3h incubation, library concentration 0,5 µM, 10 mM Mg^2+^). Candidates from band 1 of the 3rd selection round were extracted and submitted to high throughput sequencing.

We obtained ∼0.6 million readable sequences that were analysed by bioinformatics. The loop sequence of every hairpin was aligned against the reverse-complement of the loop of all other hairpins to identify potential kissing couples. the most abundant motifs in the alignments were extracted. On this basis, we identified 15 motifs of 4-7 contiguous bases in the random loop window of 93 hairpin sequences for which a complementary partner was found, i.e. that could potentially form kissing complexes.. These couples were grouped into 15 families, corresponding to such motifs and their frequencies of occurrence in round 3 (Table S2). Interestingly, the motif of loop-loop interaction in several candidate sequences (e.g. Families A1, K1, N1) includes a purine-pyrimidine palindromic DNA motif (RYRYR/YRYRY, where R and Y represents purine and pyrimidine, respectively).

For many of the tested hairpin pairs no retarded band could be detected by EMSA. This could be due to our manual selection being too wide and/or that these sequences may be low affinity candidates. Only 3 selection rounds were carried out. Further rounds of selection under more stringent conditions might have led to the elimination of these sequences.

Within the C1 family having a CCGCAC/GTGCGG consensus motif, putative DNA loop-loop complexes were identified (Table 1). Candidates were synthesized and assessed for interaction through EMSA, first in pool, then deconvoluted in individual sequences (Figure S1). An interaction was confirmed for dK25 with dK26 and dK27 (not shown), for the dK46 hairpin with dK49 and dK50, as well as dK48 with dK51, which have different flanking regions, indicating that kissing via this motif can occur in various sequence contexts. The family I1, containing the dK54/dK61 complex interacting by the AGGGGT/ACCCCT motif, gave also positive EMSA results (Table 1).

**Table 1.**
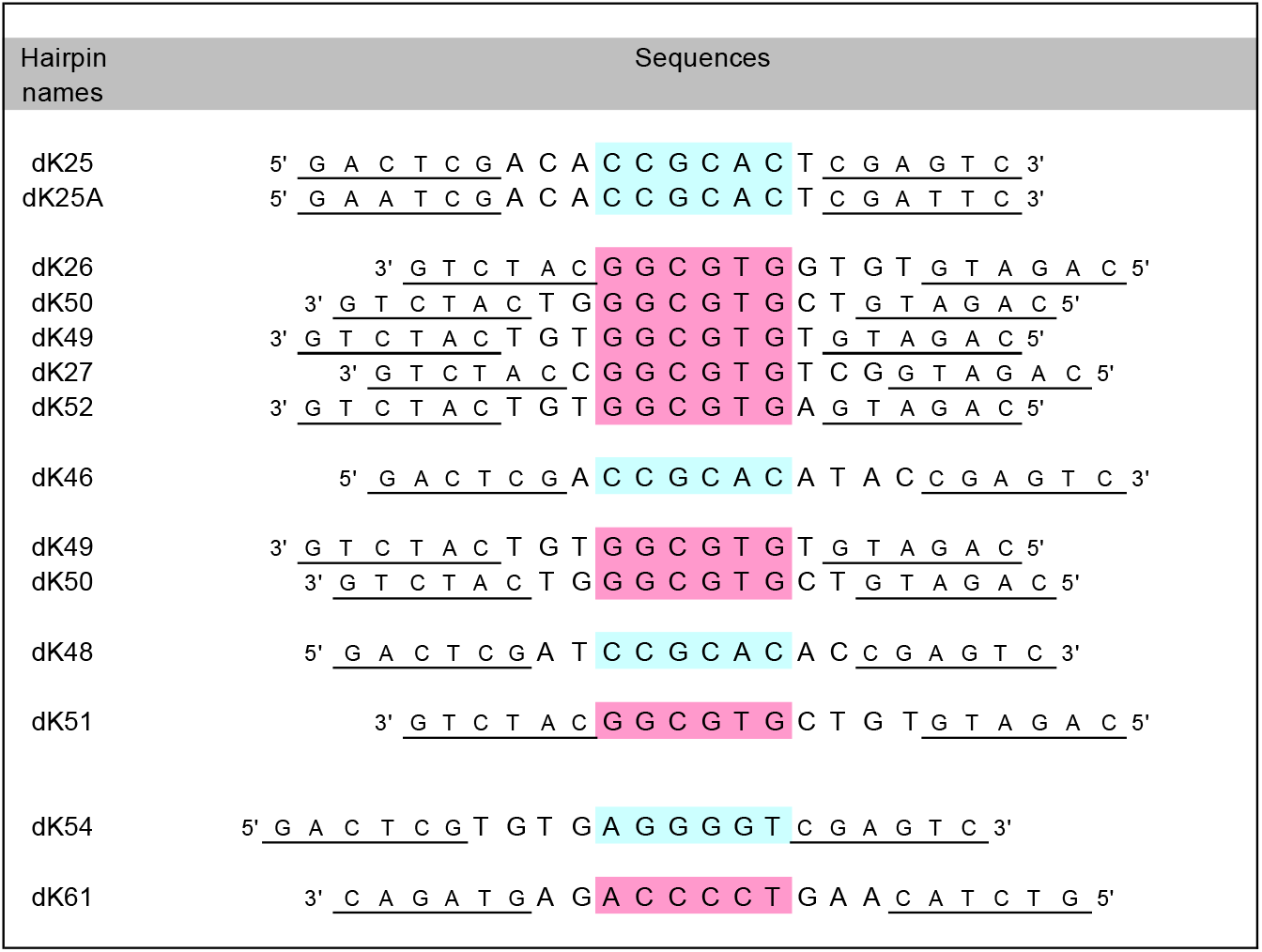
Selected hairpin oligonucleotide pairs from SELEX sequencing that can form DNA kissing loop interactions validated by EMSA. Complementary motifs that could form loop-loop interactions are in blue and pink. Underlined stem sequences are constant and complementary stems (GACTCG / CGAGTC and CAGATG / CATCTG).

The dK25-dK26 couple turned out to be the most stable complex identified. Prior to characterizing this complex, the C residue at position 3 of the stem of dK25 was replaced with A and the complementary G at position 20 with a T (resulting in sequence dK25A, Tables 1 and S1) to avoid the putative formation of extended duplexes. A putative dK25A-dK26 kissing complex structure is represented in Scheme 1a.

**Scheme 1.**
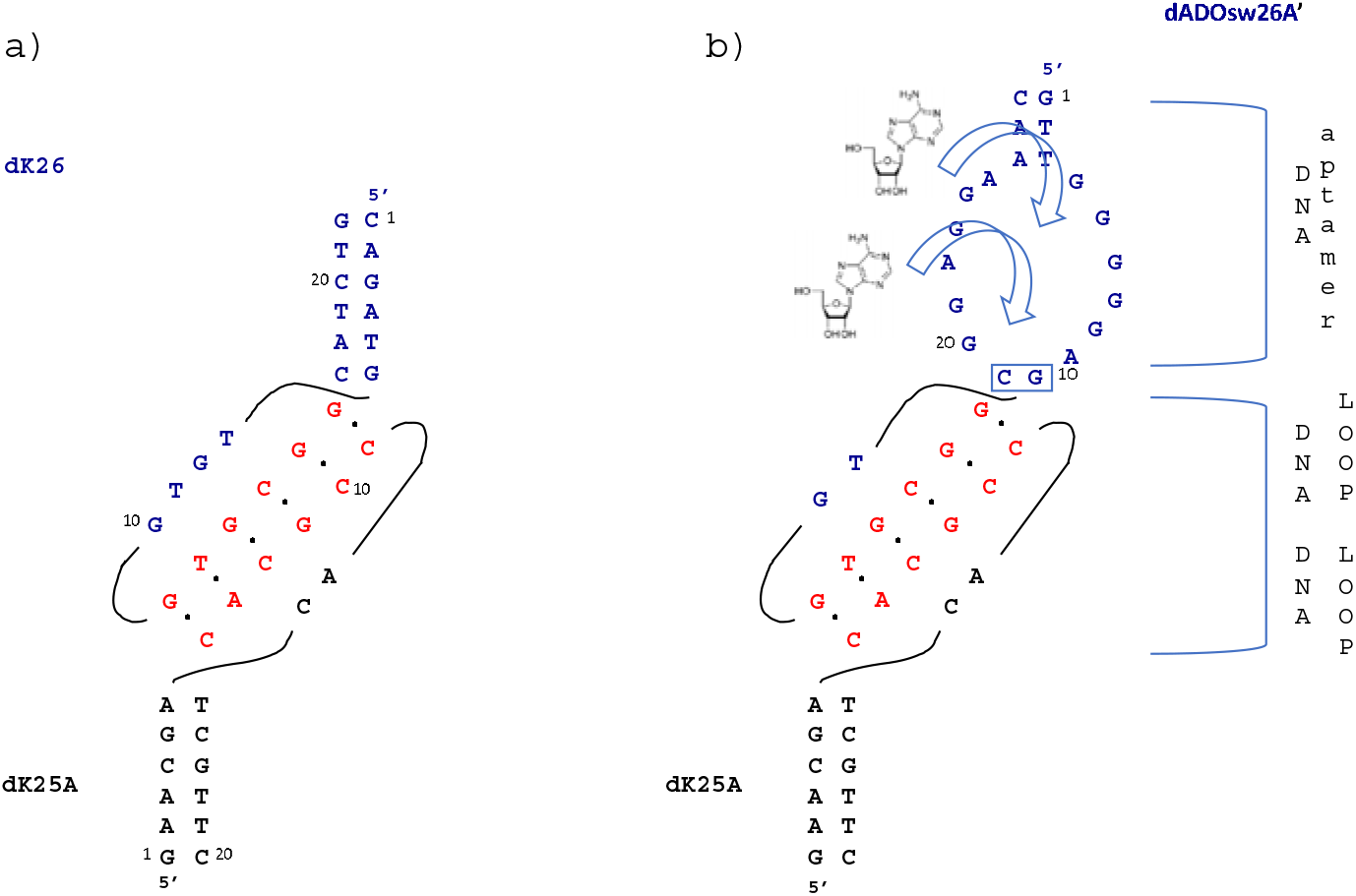
a) Schematic representation of the dK25A-dK26 kissing complex predicted from the primary sequences and b) the aptakiss-aptaswitch dK25A-dADOsw26A’ complex. dADOsw26A’ was engineered by the introduction of the dK26 loop in place of the predicted apical loop of a previously designed anti-adenosine aptamer in order to interact via a kissing complex with the loop of dK25A. The dK26 loop and the part of the aptamer that binds adenosine was linked by a G-C base pair connector.

Unfortunately, many of the tested hairpin pairs in EMSA did not exhibit retarded band. This suggested that these sequences may be low affinity candidates. Further rounds of selection under more stringent conditions might discard these low affinity sequences.

### Characterization of a DNA-DNA kissing complex

Since kissing complexes rely on high Mg concentration, the affinity of dK25A-dK26 complex in the presence of 0, 10 or 50 mM of magnesium acetate was determined by fluorescence anisotropy (FA). As expected, KD values increased from 160 ± 50 nM at 0 mM Mg2+, to 55 ± 6 nM at 10 mM Mg2+ and 41 ± 12 nM at 50 mM Mg2+. We next investigated the KD of the Mg2+ for the complex, which was evaluated at 11.5 +/-1.5 mM (Fig 2d). In addition, the specificity of the loop-loop interaction was confirmed by FA using a dK26 A13 mutant (Table S1). The G13 nucleotide located in the centre of the kissing motif of the dK26 loop (scheme 1) was switched to A in order to abolish the loop-loop interaction with dK25A. (data not shown) confirming that the two partners interact through their loops and that G13 is essential to the recognition motif.

**Figure 2.**
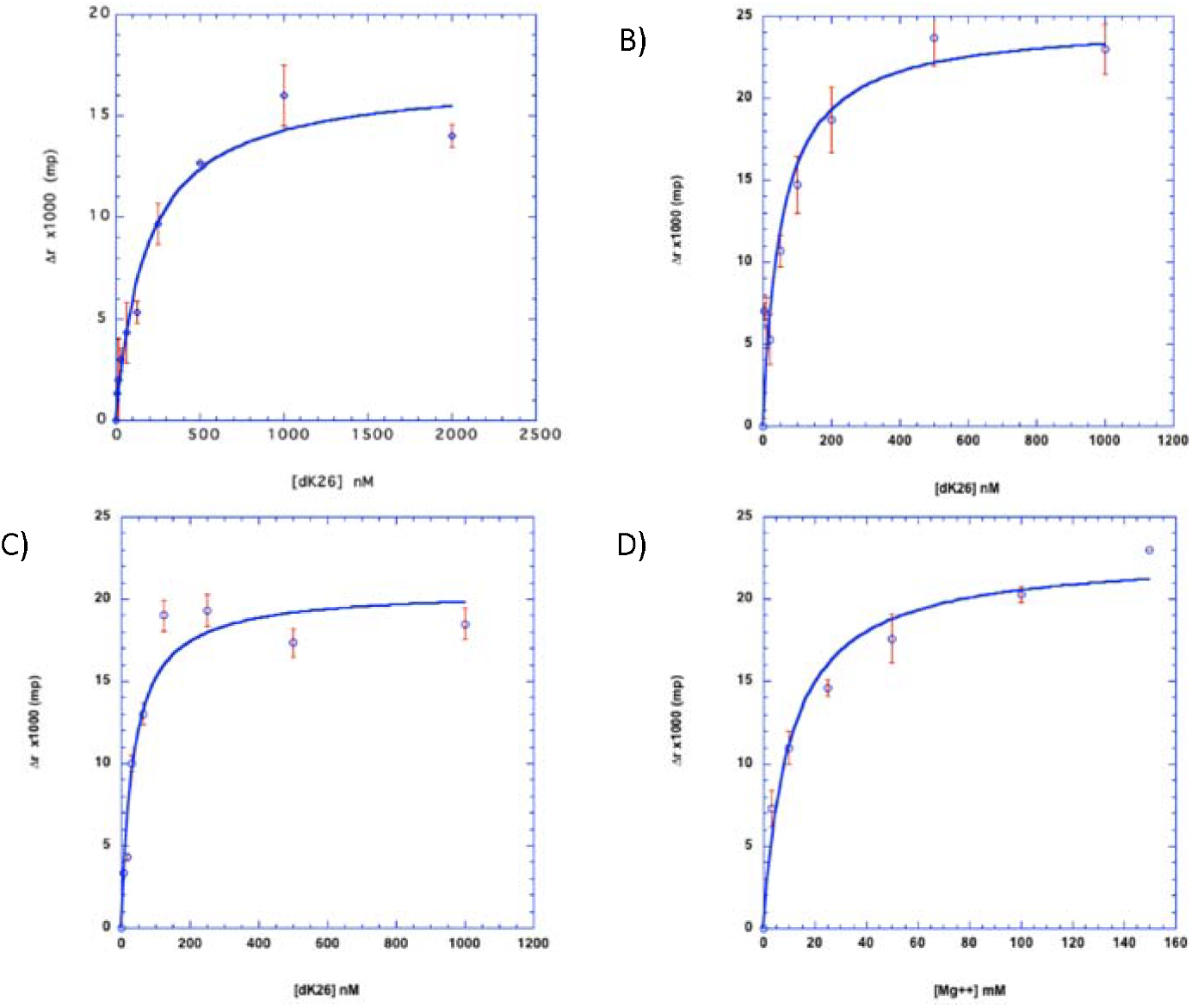
Characterisation of the dK25A-dK26 complex by fluorescence anisotropy (FA). Binding affinity of the dK25A-dK26 kissing complex at increasing concentrations of magnesium : a) 0 mM b) 10 mM and c) 50 mM. [Rox-dk25A] = 10 nM. d) Binding affinity of the Mg2+ for the selected dK25A-dK26 complex. [Rox-dk25A] = 10 nM ; [dk26] = 1 µM.. Experiments were performed 2 or 3 times in triplicate, and for each KD determinations, the result of one experiment is represented.

### Design and characterization of a full DNA aptakiss-adenoswitch

RNA and DNA adenoswitches have been previously designed to bind to RNA aptakiss partners via the formation of RNA-RNA or RNA-DNA kissing complexes, specific to the binding of adenosine^[15, 23]^. Based on these previous designs, we developed aptaswitches using the DNA kissing complex dK25A-dK26 (table 2). A predicted apical loop of the DNA aptamer directed against the adenosine which is not involved in the binding of its target molecule was replaced by the DNA loop sequence of the dK26. The DNA hairpin dK25A whose loop is partially complementary of the DNA loop of dK26 constituted the DNA aptakiss. As it has been determined that nucleic acid bases located at the loop-aptamer junction were critical to the stability of the full RNA aptakiss-aptaswitch complexes^[1]^, different lengths and sequences of these connecting positions were designed and their affinity evaluated through Fluorescence Anisotropy variation measurement (FA) . Two adenoswitches dADOsw26A’ and dADOsw26B’ were engineered, composed of two loops : an internal one for adenosine binding, an apical one for kissing interaction, connected with one G/C (dADOsw26A’) or two CA/TG base pairs (TG, dADOsw26B’). The link between the dK26 stem and the motif of interaction with dK25A composed of TGTG was reduced to one TG. In an opposite strategy, adenoswitches were designed from the dK25A loop in place of the dK26 loop, containing in the dADOsw26A’/B’ (Table 2). In this case, dK26 became the aptakiss. Three adenoswitches (dADOsw25A’, dADOsw25B’ and dADOsw25C’) showing three different sequences at their loop-aptamer junctions (GA/TC ; CA/TG and A/T) were designed (Table 2). The connector between the loop and the adenosine aptamer region was designed of maximum 3 base pairs. Furthermore, the stem of the parent aptamer was truncated at 3 base pairs as previously described for RNA/DNA-complexes^[22a]^.

**Table 2.**
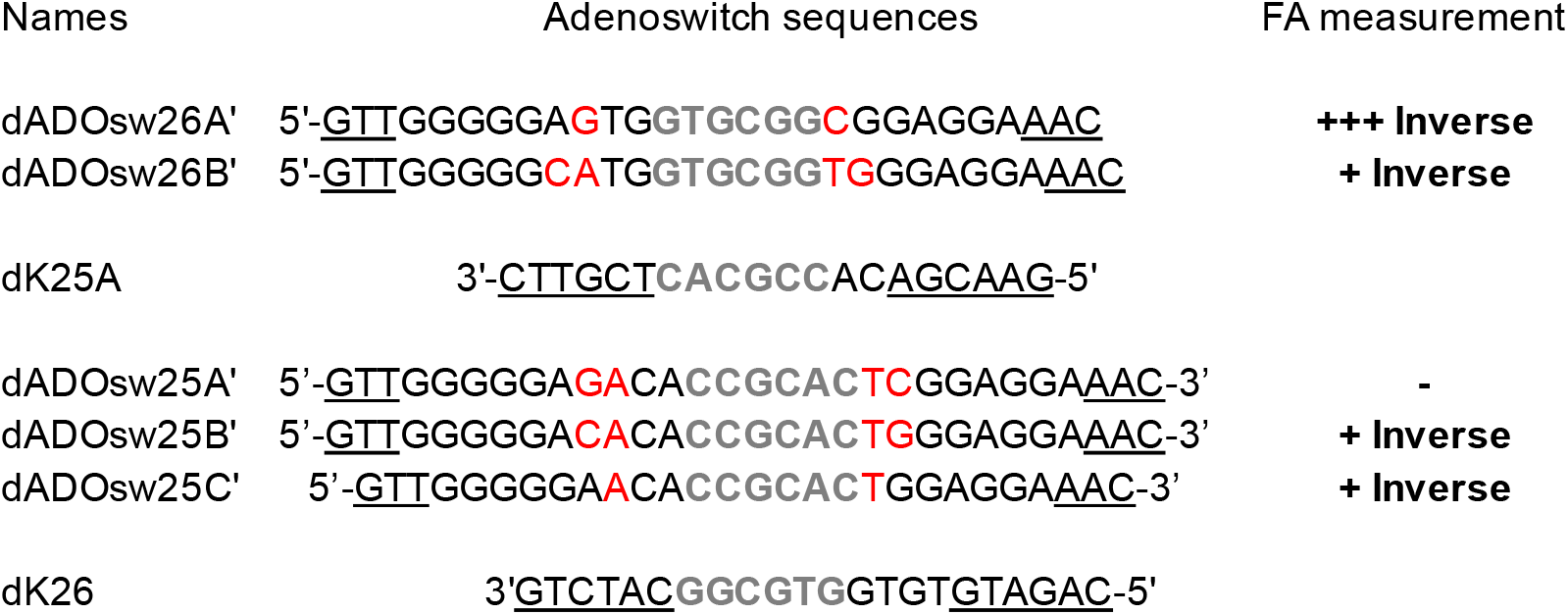
Sequences of the designed DNA-adenoswitches. The loop-loop motifs of interaction are in grey. Connectors are indicated in red. The aptakiss-aptaswitch interactions estimated by FA are given on the right and are described in the text.

These five full DNA adenoswitches were tested in solution by FA for their ability to form complexes with the fluorescent-labeled Rox-dK25A or TR-dK26 aptakisses in the presence of adenosine (Table 2). The FA measurements show a decreased intensity with the addition of adenosine for most of the complexes reflecting the dissociation of the complex (data not shown). The highest decrease of FA intensity was observed when Rox-dK25A was mixed with the adenoswitch dADOsw26A’. Thus, the study was further focused on this aptaswitch complex dK25A-dADOsw26A’ (Scheme 1b). The dADOsw26A’ switch binds to dK25A aptakiss with an K_D_ of 115 nM ± 5 nM (Figure S3), in the R buffer containing 10 mM of magnesium.

**Scheme 1.**
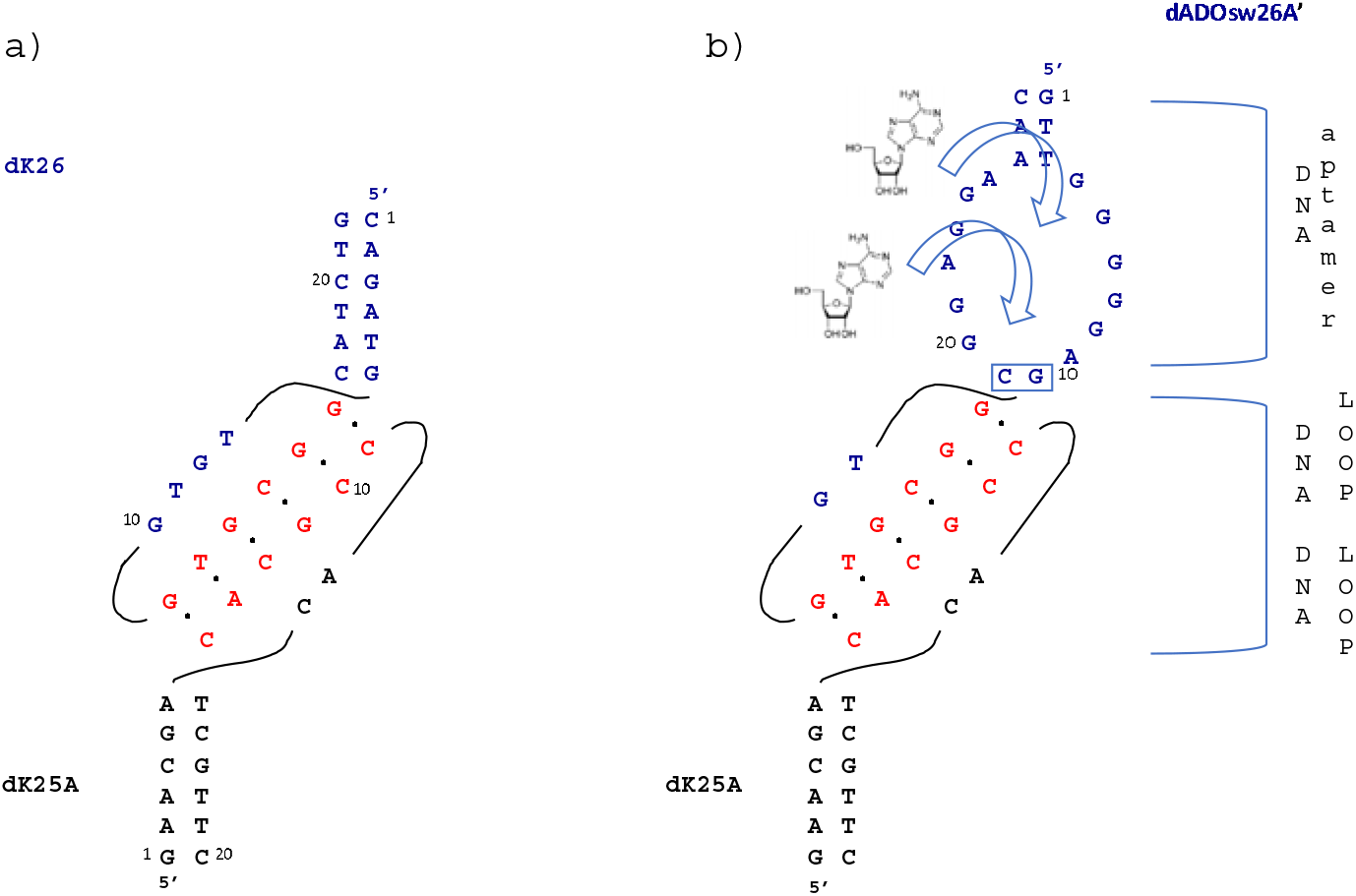
a) Schematic representation of the dK25A-dK26 kissing complex predicted from the primary sequences and b) the aptakiss-aptaswitch dK25A-dADOsw26A’ complex. dADOsw26A’ was engineered by the introduction of the dK26 loop in place of the predicted apical loop of a previously designed anti-adenosine aptamer in order to interact via a kissing complex with the loop of dK25A. The dK26 loop and the part of the aptamer that binds adenosine was linked by a G-C base pair connector.

In order to build an aptasensor, the affinity of this complex for adenosine was determined by FA (Figure 4). Results showed that adenosine induced a conformational change of the dADOsw26A’ resulting in a dissociation of the dK25A-dADOsw26A’ complex. The apparent dissociation constant of adenosine for the dK25A-dADOsw26A’ couple was estimated by FA at 700 +/-100 µM (Figure 4a). As compared to the parent DNA aptamer (K_D_ = 6 +/-3 µM^[26]^), a factor 100 in affinity for adenosine was lost . The specificity was investigated at different concentrations of Mg^2+^, (i.e., 0, 10 and 50 mM), by adding 2 mM of inosine, a small molecule which differs from adenosine only by an amine groupment and that doesn’t bind to the anti-adenosine aptamer (Fig 4b). Under these conditions, only low signals of fluorescence were detectable,, which demonstrated the specificity of dK25A-dADOsw26A’ complex to adenosine.

Finally, we checked the specificity of dADOsw26A’ by mixing a mutant of the adenosine recognition loop (A15) (Table S1) and dK25A. We observed that the sensitivity to adenosine was kept at a concentration higher than 1 mM with the mutated dADOsw26A’(A6-7) sequence (Table S1) corresponding to an aptamer which is unable to bind adenosine.

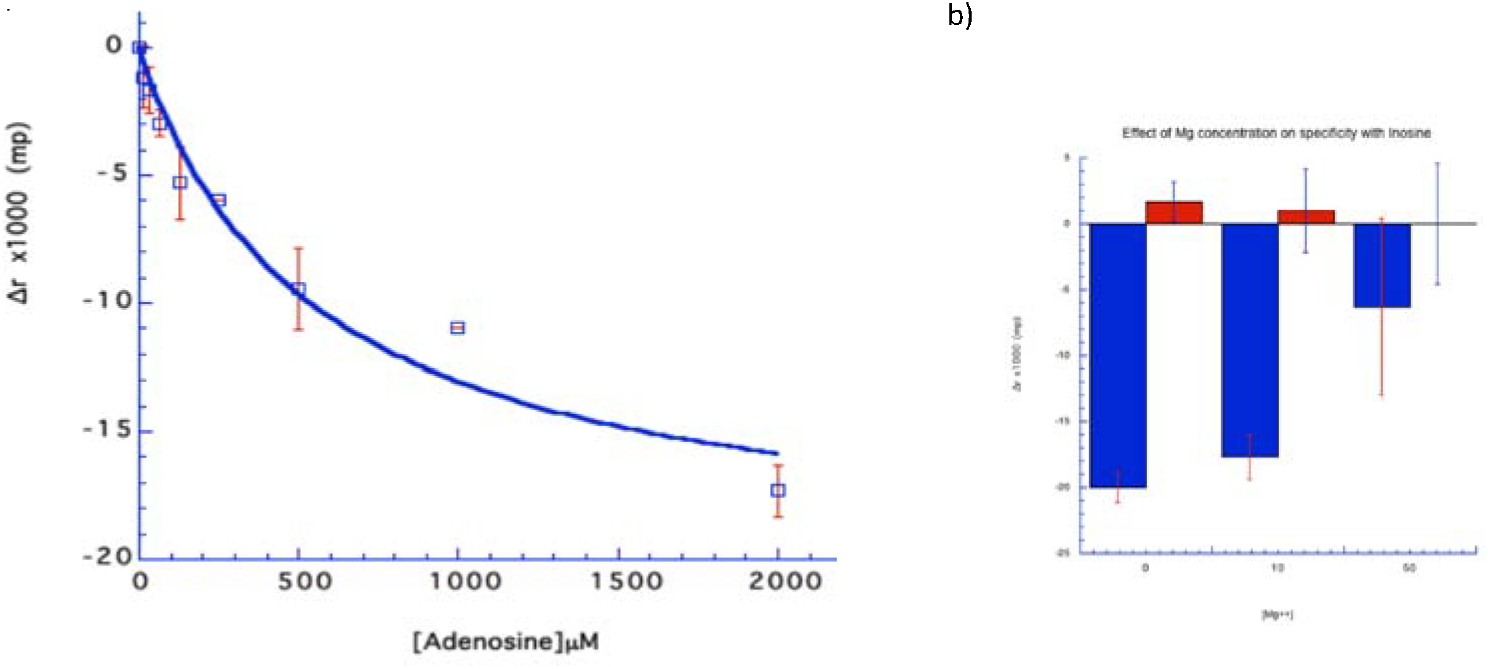

### Experimental

#### DNA Libraries and oligonucleotides

Oligonucleotide sequences used in this study are reported in Table S1. The primer P3 was labelled at the 5’end with a biotin and used to produce single-stranded DNA by asymmetric PCR. All oligonucleotides used in this study were purified by electrophoresis on denaturing 20% polyacrylamide, 7M urea gels.

#### In vitro selection (SELEX)

Before each round of selection, DNA pools were structured by heating at 90°C for 2 min in water, chilled on ice for 2 min and completed with the R selection buffer (final concentration: 20 mM HEPES, 20 mM sodium acetate, 140 mM potassium acetate, and 3 mM magnesium acetate, pH 7.4) at room temperature. The library (100 picomoles) was mixed in a final volume of 20 µL for 12 h. In the first round of selection, the stringency was low enough to select a pool of DNA sequences able to form kissing complexes whatever their affinities. In subsequent rounds, to keep only high stability complexes, the DNA hairpin concentration was decreased 10 times at each round. Moreover, the time of incubation was decreased (6 h for round 2 and 3 h for the final round). DNA population was separated by EMSA. Samples were run on a native gel (15% [w/v], 75:1 acrylamide:bis-acrylamide) in 50 mM Tris-acetate (pH 7.3 at 20°C) and 3 mM magnesium acetate (TAC buffer) at 100 V and 4°C for 15 h). The gels were stained 20 min with 0.5 µg/mL of ethidium bromide and complexes were visualized by UV. The shifted bands corresponding to DNA complexes were extracted from the gel, eluted for 1 h at 65°C in 500 µL of the elution buffer [10 mM Tris-HCl, pH 7.4, 1 mM EDTA (EthyleneDiamineTetracetic Acid), and 25 mM NaCl], and then, ethanol precipitated. Round 2 was performed without magnesium acetate to avoid selection of DNA duplexes. For this round of SELEX, free DNA hairpins that were not engaged in complexes were purified and used for the next round of selection (Fig S1).

#### DNA amplification and sequencing of the candidates

Purified DNA hairpins were amplified using 25 units of AmpliTaq Gold polymerase (PE Applied Biosystems) in 500 µL of the Taq polymerase buffer containing 200 µM of dNTP, 1.5 mM of Mg2+ and 5% of DMSO. Single-stranded products were synthetised by asymmetric PCR using 2 µM of P5 primer and 0.2 µM of the biotinylated P3 primer. Then, the reaction mixture was denatured at 94°C for 10 min to activate the enzyme and was subjected to repeated cycles: 94°C for 1 min, 60°C for 1 min, 72°C for 1 min 30 sec for 20 cycles and 72°C for 10 min, for one final cycle. PCR products were purified with a 10K Nanosep device (PALL©). The biotinylated double-stranded PCR product was eliminated using streptavidine MagneSphere© paramagnetic coated beads (Promega©). Single-stranded DNA remaining in the supernatant was quantified by absorbance at 260 nm and used for the next round. After 3 rounds of selection, DNA candidates were sequenced by Next Generation Sequencing (Illumina^®^ technology, platform of the “Institut du Cerveau et de la Moëlle”, Paris).

#### Bioinformatic sequence analysis

DNA from round 3 was sequenced at Integragen, Inc. (Évry, France) following the Illumina short-read approach based on 37-nt single-end reads. About 600,000 sequences were obtained. Paired reads were assembled into a single sequence using PANDAseq 2.9^[27]^ run with options “-o 10 -O 150 -t 0.6 -A simple_bayesian -C empty”. Assembled sequences that contained hairpins with stem regions identical to the designed sequences (5’-GACTCG-(12 nt)-CGAGTC-3’ and 5’-CAGATG-(8 nt)-CATCTG-3’) were retained. Assembled sequences may contain both or either one of these hairpins. The number of retained sequences ∼330,000. In order to identify kissing motifs, the loop regions of the sequences were compared. Sequences were folded using Mfold 3.6^[28]^ and loop sub sequences were extracted based on the structure information from the “.det” output file of Mfold. The folding parameters of Mfold were set to the values used during the SELEX process, i.e., for round 3, a Na+ concentration (NA_CONC parameter) of 0.020M, a Mg2+ concentration (MG_CONC) of 0.003M, and at room temperature (T). Extracted loop regions ranged from 3 to 12 nucleotides in length. For sequences that contained a single loop, the loop region of every sequence was aligned against the reverse-complement of the loop region of every other sequence. Optimal global alignments were computed using the Needleman and Wunsch algorithm as implemented in the “needle” program (run with options “-gapopen 10 - gapextend 0.5”) of the EMBOSS 6.6.0 package^[29]^. The “GNU parallel” utility[30] was used to dispatch large numbers of alignments to multiple processors. Then, the alignments were parsed in order to extract those that contained contiguous stretches of 3 or more matching nucleotides. The most abundant motifs observed in the alignments were selected for experimental validation as putative kissing motif candidates The choice was based on whether the frequency was substantially higher that of the other motifs, and whether the motif was found among top motifs in multiple analyses.

#### Fluorescence anisotropy (FA) measurements

Prior to be used, samples were treated in the same way as were the libraries during the selection. The dK25A hairpin labelled with a Rox (Rhodamine X) group at its 5’end was used at 10 nM in a final volume of 50 µL of R buffer. Complex with dk26 or aptaswitches were incubated during 2 hours at 4°C in the dark. Experiments were carried out two times in triplicate at either 0 or 10 or 50 mM of Mg^2+^ concentration, on a TECan^®^ Ultimate 3000 platereader, in Fluorescence Polarization measurement mode (Ex. 590 nm, Em 620 nm).

## Conclusion

In this study we reported the selection and characterization of de novo DNA/DNA kissing complexes from an in vitro selection process. Four kissing motifs were identified and confirmed by EMSA. Among them, the couple dk25/dk26 exhibited a nanomolar affinity (K_D_ = 41 ± 12 nM with 50 mM Mg^2+^) revealed by fluorescence anisotropy, relying on magnesium, which confirmed the loop-loop interaction. Such kissing candidates were integrated in the design of new aptaswitch-aptakiss biosensors only made of DNA that respond to the presence of adenosine. From the four aptakiss/aptaswitch candidates, dK25A-dADOsw26A’ complex exhibited a conformational change that dissociates the kissing complex in the presence of adenosine, but not inosine.

This first report of a DNA/DNA aptaswitch/aptakiss complex could be used to build nanotechnology system with conditional interaction upon a small molecule.

## Conflicts of interest

There are no conflicts to declare.

## Notes

### Competing Interest Statement

The authors have declared no competing interest.

## Notes and references

[1] F. Duconge, C. Di Primo, J. J. Toulme, J. Biol. Chem. 2000, 275, 21287–21294.

[2] K. Y. Chang, I. Tinoco, Proc. Natl. Acad. Sci. USA 1994, 91, 8705–8709; bE. V. Pilipenko, S. V. Maslova, A. N. Sinyakov, V. I. Agol, Nucleic Acids Res. 1992, 20, 1739–1745.

[3] X. Ling, D. Golovenko, J. Gan, J. Ma, A. A. Korostelev, W. Fang, Nature 2025, 644, 1107–1115.

[4] C. Brunel, R. Marquet, P. Romby, C. Ehresmann, Biochimie 2002, 84, 925–944.

[5] Y. Eguchi, J.-i. Tomizawa, Cell 1990, 60, 199–209.

[6] aF. Jossinet, J. C. Paillart, E. Westhof, T. Hermann, E. Skripkin, J. S. Lodmell, C. Ehresmann, B. Ehresmann, R. Marquet, RNA (New York, N.Y.) 1999, 5, 1222–1234; bN. Windbichler, M. Werner, R. Schroeder, Nucleic Acids Res. 2003, 31, 6419–6427.

[7] L. Argaman, S. Altuvia, Journal of Molecular Biology 2000, 300, 1101–1112.

[8] H. Arnion, Dursun N. Korkut, S. Masachis Gelo, S. Chabas, J. Reignier, I. Iost, F. Darfeuille, Nucleic Acids Res. 2017, 45, 4782–4795.

[9] aS. Shetty, S. Stefanovic, M. R. Mihailescu, Nucleic Acids Res. 2013, 41, 2526–2540; bC. Romero-López, A. Berzal-Herranz, RNA 2009, 15, 1740–1752; cP. Friebe, J. Boudet, J.-P. Simorre, R. Bartenschlager, J. Virol. 2005, 79, 380–392; dE. Ennifar, J.-C. Paillart, A. Bodlenner, P. Walter, J.-M. Weibel, A.-M. Aubertin, P. Pale, P. Dumas, R. Marquet, Nucleic Acids Res. 2006, 34, 2328–2339; eA. H. Kensinger, J. A. Makowski, M. R. Mihailescu, J. D. Evanseck, ACS Phys. Chem. Au 2025, 5, 410–424.

[10] aI. Severcan, C. Geary, A. Chworos, N. Voss, E. Jacovetty, L. Jaeger, Nat. Chem. 2010, 2, 772–779; bC. Geary, A. Chworos, E. Verzemnieks, N. R. Voss, L. Jaeger, Nano Lett. 2017, 17, 7095–7101; cI. Severcan, C. Geary, E. Verzemnieks, A. Chworos, L. Jaeger, Nano Lett. 2009, 9, 1270–1277; dS. Horiya, X. Li, G. Kawai, R. Saito, A. Katoh, K. Kobayashi, K. Harada, Chem. Biol. 2003, 10, 645–654.

[11] aH. Udono, M. Fan, Y. Saito, H. Ohno, S.-i. M. Nomura, Y. Shimizu, H. Saito, M. Takinoue, ACS Nano 2024, 18, 15477–15486; bAnli A. Tang, Martin V. Gobry, S. Li, Ebbe S. Andersen, E. Franco, Nucleic Acids Research 2025, 53, gkaf497.

[12] M. Xie, W. Chen, M. Vonk-de Roy, A. Walther, Nature Communications 2025, 16, 8254.

[13] F. Ducongé, J. J. Toulmé, RNA (New York, N.Y.) 1999, 5, 1605–1614.

[14] aM. Watrin, F. Von Pelchrzim, E. Dausse, R. Schroeder, J.-J. Toulmé, Biochemistry 2009, 48, 6278–6284; bJ. J. Toulmé, C. Di Primo, S. Moreau, in Progress in Nucleic Acid Research and Molecular Biology, Vol. 69, Academic Press, 2001, pp. 1–46.

[15] G. Durand, E. Dausse, E. Goux, E. Fiore, E. Peyrin, C. Ravelet, J.-J. Toulmé, Nucleic Acids Res. 2016, 44, 4450–4459.

[16] aC. Boiziau, E. Dausse, L. Yurchenko, J.-J. Toulmé, J. Biol. Chem. 1999, 274, 12730–12737; bD. Sekkai, E. Dausse, C. Di Primo, F. Darfeuille, C. Boiziau, J. J. Toulme, Antisense Nucleic Acid Drug Dev 2002, 12, 265–274; cF. Darfeuille, D. Sekkai, E. Dausse, G. Kolb, L. Yurchenko, C. Boiziau, J. J. Toulmé, Comb. Chem. High Throughput Screen. 2002, 5, 313–325.

[17] aR. Nutiu, Y. Li, Angew. Chem. Int. Ed. 2005, 44, 1061–1065; bR. Stoltenburg, N. Nikolaus, B. Strehlitz, Journal of Analytical Methods in Chemistry 2012, 2012, 1–14; cK. A. Yang, R. Pei, M. N. Stojanovic, Vol. 106, Academic Press, 2016, pp. 58–65; dK.-A. Yang, M. Barbu, M. Halim, P. Pallavi, B. Kim, D. M. Kolpashchikov, S. Pecic, S. Taylor, T. S. Worgall, M. N. Stojanovic, Nat. Chem. 2014, 6, 1003–1008; eK. Yang, N. M. Mitchell, S. Banerjee, Z. Cheng, S. Taylor, A. M. Kostic, I. Wong, S. Sajjath, Y. Zhang, J. Stevens, S. Mohan, D. W. Landry, T. S. Worgall, A. M. Andrews, M. N. Stojanovic, Science 2023, 380, 942–948.

[18] T. A. Feagin, N. Maganzini, H. T. Soh, ACS Sensors 2018, 3, 1611–1615.

[19] aE. Goux, E. Dausse, V. Guieu, L. Azéma, G. Durand, M. Henry, L. Choisnard, J. J. Toulmé, C. Ravelet, E. Peyrin, Nanoscale 2017, 83, 6464–6467; bM. N. Stojanovic, P. de Prada, D. W. Landry, J. Am. Chem. Soc. 2001, 123, 4928–4931; cR. Nutiu, Y. Li, J. Am. Chem. Soc. 2003, 125, 4771–4778.

[20] J. Canoura, Y. Liu, O. Alkhamis, Y. Xiao, Analytical Chemistry 2023, 95, 18258–18267.

[21] aS. Dalirirad, D. Han, A. J. Steckl, ACS Omega 2020, 5, 32890–32898; bB. Chovelon, V. Ranganathan, S. Srinivasan, E. M. McConnell, P. Faure, E. Fiore, C. Ravelet, E. Peyrin, M. DeRosa, Anal. Chem. 2024, 96, 6875–6880.

[22] aG. Durand, S. Lisi, C. Ravelet, E. Dausse, E. Peyrin, J.-J. Toulmé, Angew. Chem. Intl. Ed. 2014, 53, 6942–6945; bA. Sett, L. Zara, E. Dausse, J. J. Toulmé, Talanta 2021, 232, 122417.

[23] L. Azéma, S. Bonnet-Salomon, M. Endo, Y. Takeuchi, G. Durand, T. Emura, K. Hidaka, E. Dausse, H. Sugiyama, J.-J. Toulmé, Nucleic Acids Res. 2018, 46, 1052–1058.

[24] A. Barth, D. Kobbe, M. Focke, Nucleic Acids Res. 2016, 44, 1502–1513.

[25] K. I. Takahashi, S. Baba, P. Chattopadhyay, Y. Koyanagi, N. Yamamoto, H. Takaku, G. Kawai, RNA 2000, 6, 96–102.

[26] D. E. Huizenga, J. W. Szostak, Biochemistry 1995, 34, 656–665.

[27] A. P. Masella, A. K. Bartram, J. M. Truszkowski, D. G. Brown, J. D. Neufeld, BMC Bioinformatics 2012, 13, 31.

[28] M. Zuker, Nucleic Acids Res 2003, 31, 3406–3415.

[29] P. Rice, I. Longden, A. Bleasby, Trends in Genetics 2000, 16, 276–277.

[30] O. Tange, The USENIX Magazine 2011, 42–47.

