## supplementary data for "Aptaswitch/aptakiss adenosine sensor design based on the selection of new DNA/DNA kissing complexes"

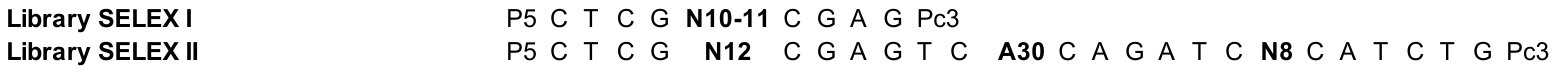

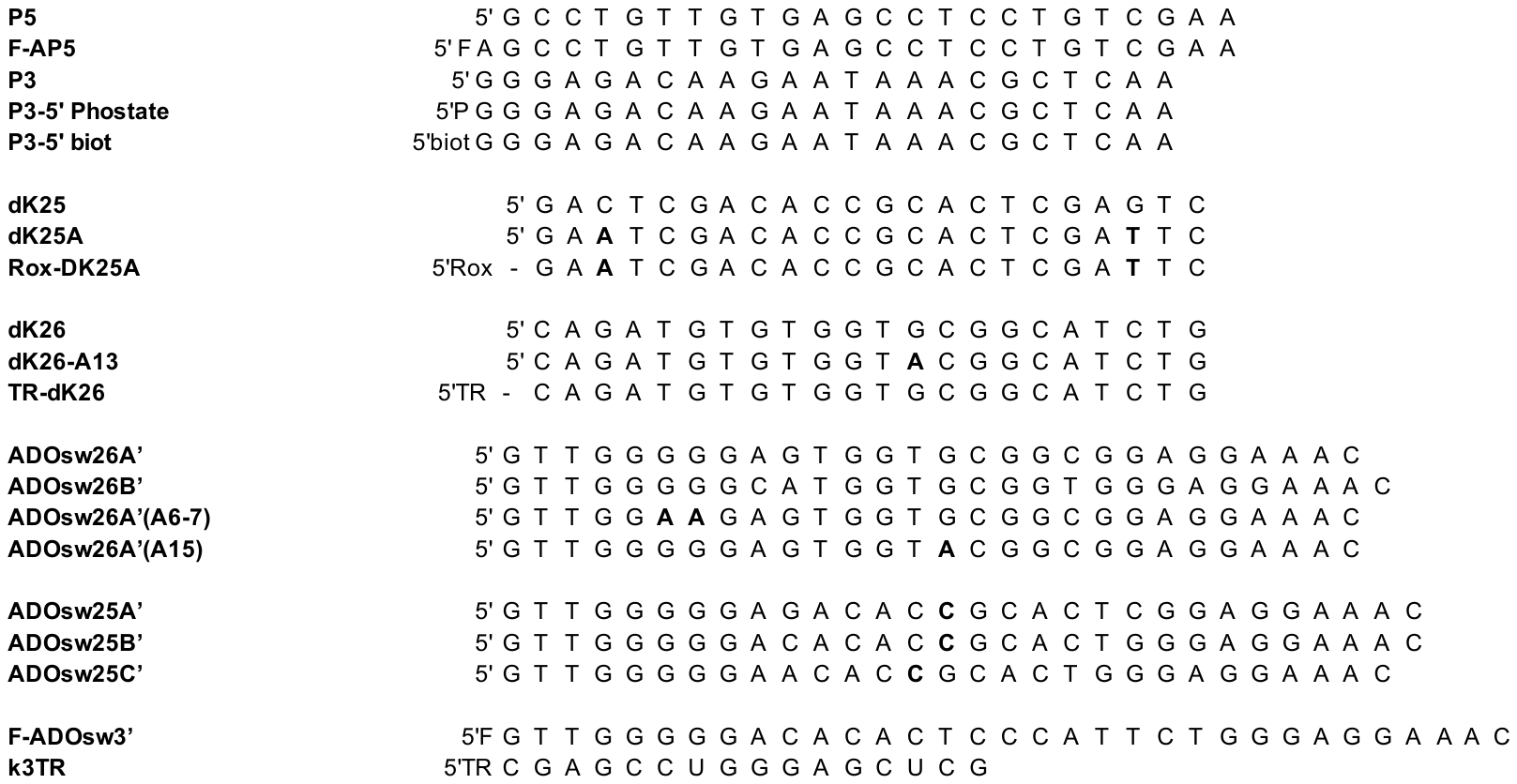

Table S1. Sequences used in this study are reported from 5’ to 3’. Abbreviations: P, phosphate; biot, biotin; Rox, carboxy-X-rhodamine; TR, Texas Red; F, Fluorescein; Pc3, complementary sequence of P3. Mutations are indicated in bold.

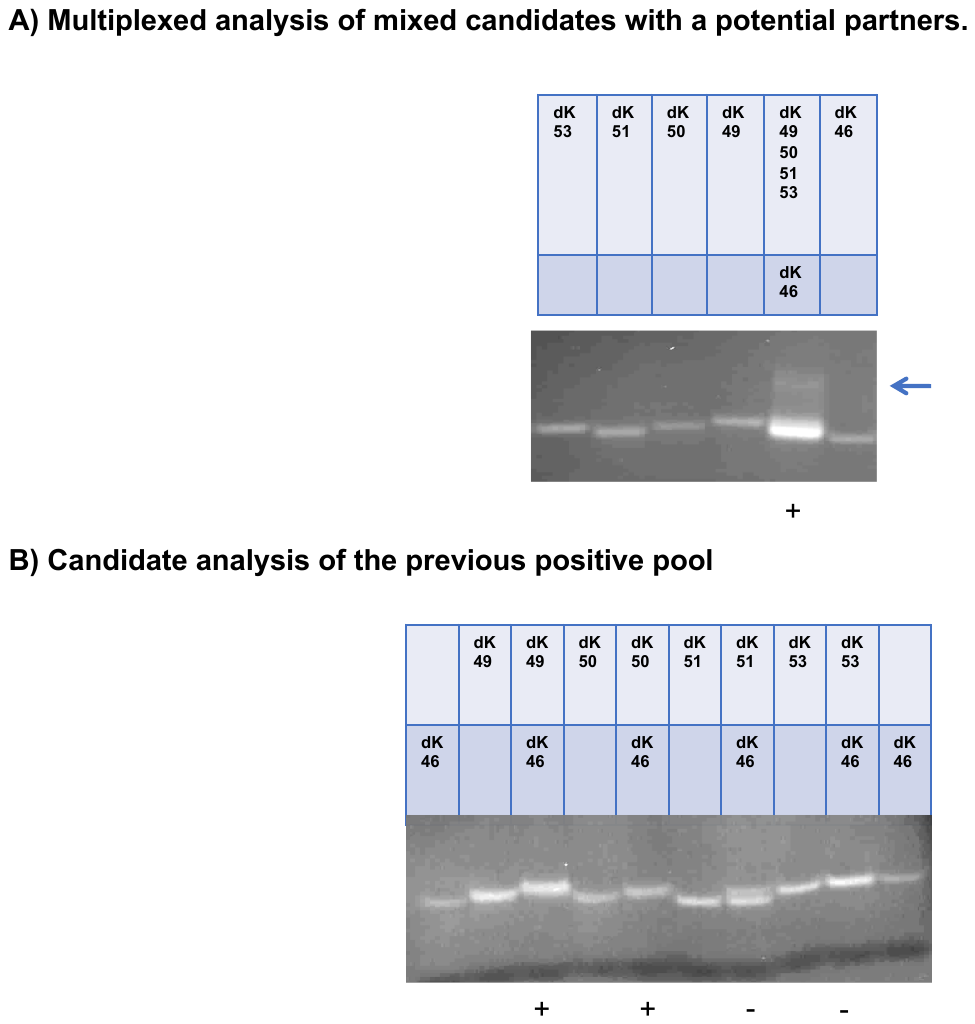

Fig S1) **Native gel electrophoresis of putative kissing complexes.** Kissing complex formation were checked by Electrophoretic Mobility Shift Assay (EMSA) on 15% polyacrylamide gels. Hairpins were incubated at 1 μM with 1 µM of putative partner at 4°C. A) Candidates of a selected family (C1b; see Table S2) were multiplexed. Candidates dK49, dK50, dK51, dK53 were pooled and mixed with the putative hairpin partner dK46, and each free hairpin was also run separately as a migration control. Positive results showing a smear (blue arrow on the right of the gel) when the mixture was run and colored with ethidium bromide were deconvoluted. B) Deconvolution analysis of the positive pool. Each hairpin of the positive mixture was tested by EMSA against its predicted hairpin partner, i.e., dK46 with dK49, dK46 with dK50, dK46 with dK51, and dK46 with dK53. A lower migration than expected for the free hairpins may indicate the formation of a putative complex (see main text for details).

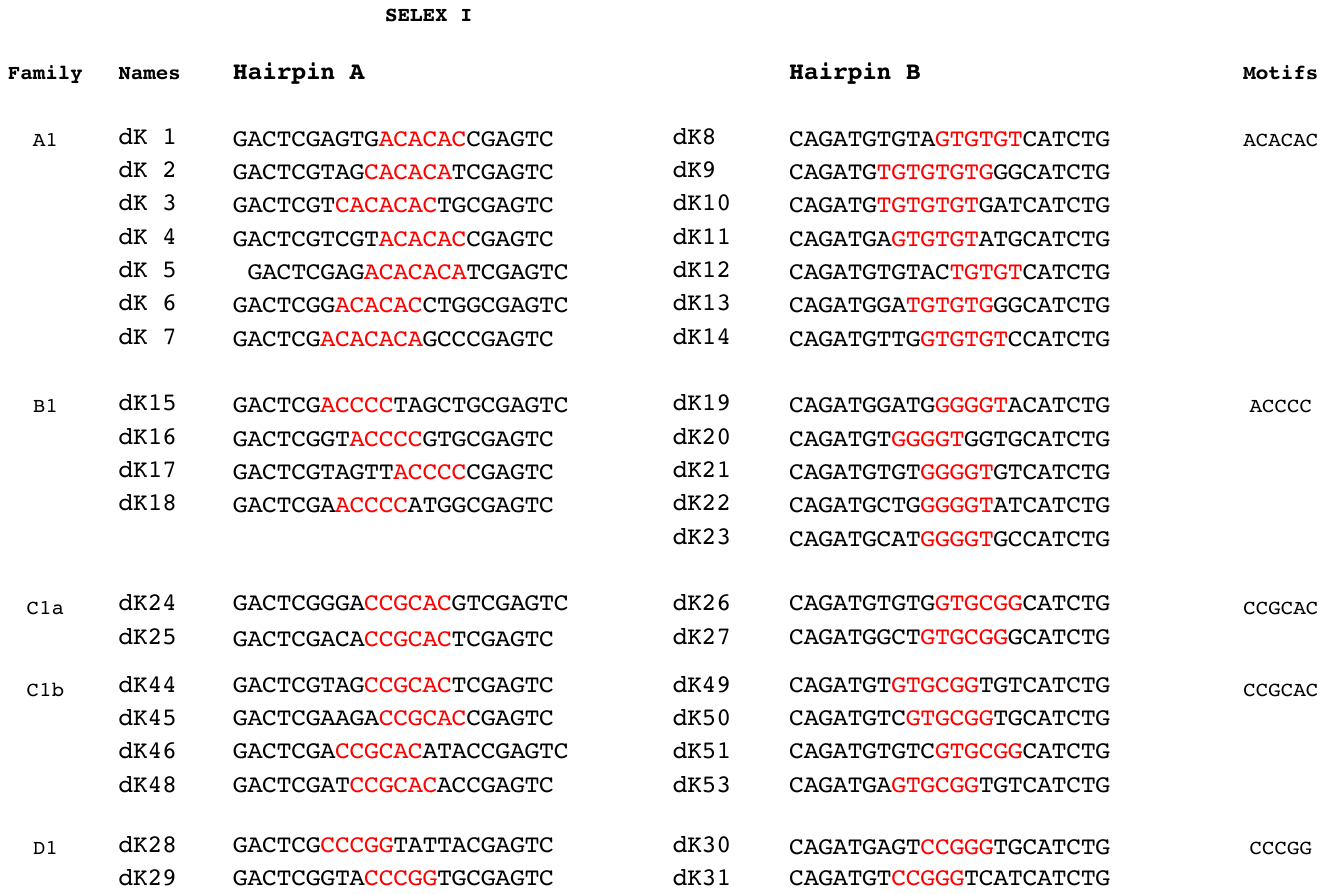

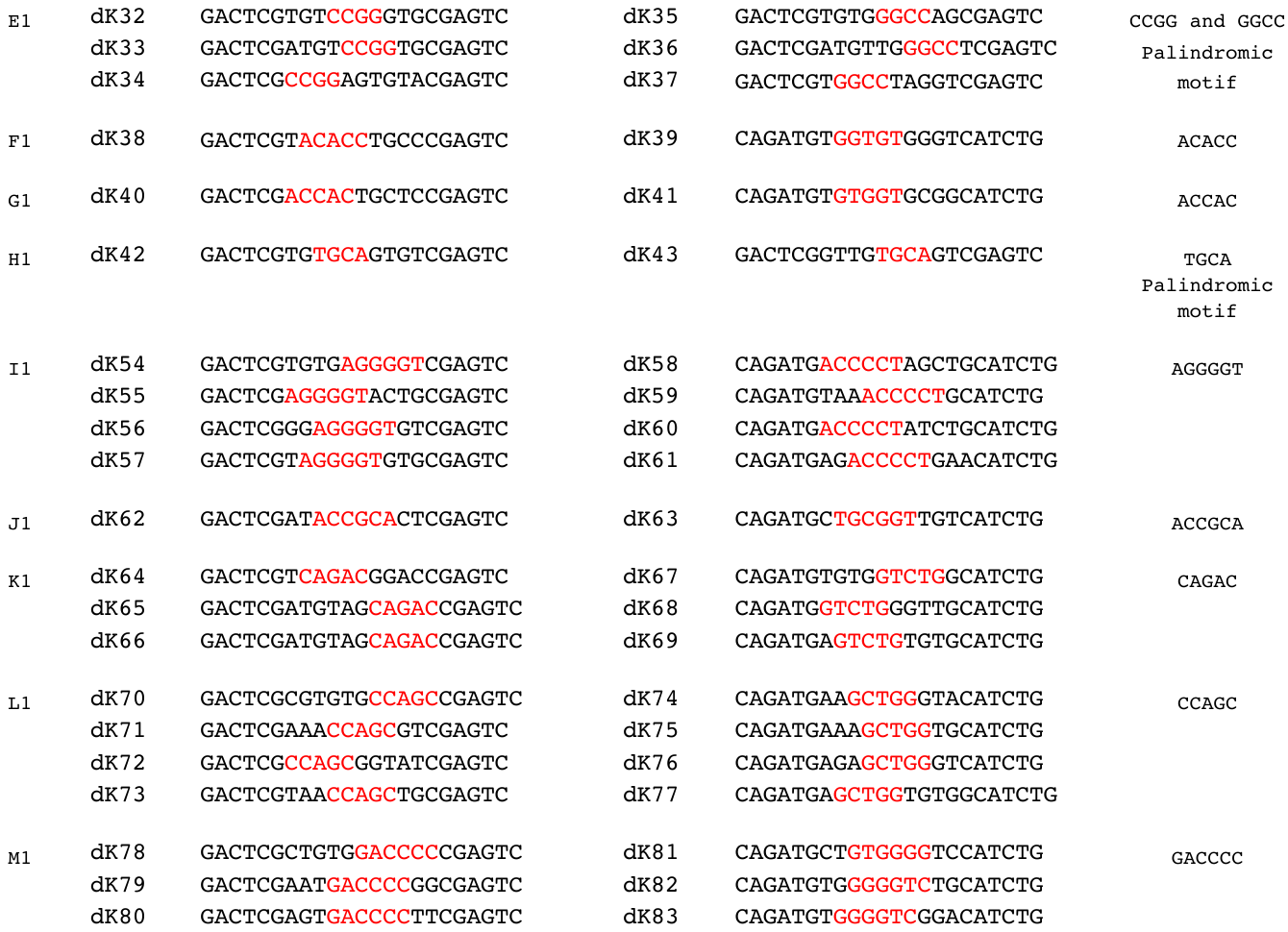

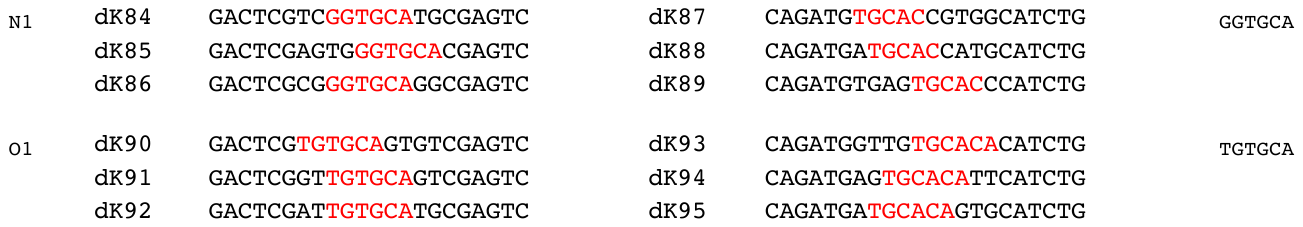

**Table S2)** Sequences selected from bioinformatic analysis which have been synthesized and checked for loop-loop interaction.

Complementary kissing motifs are indicated in red. Sequences were grouped into families based on the motif indicated in the rightmost column. They are listed from 5’ to 3’ from left to right. Some sequences may appear several times under different names and motifs.

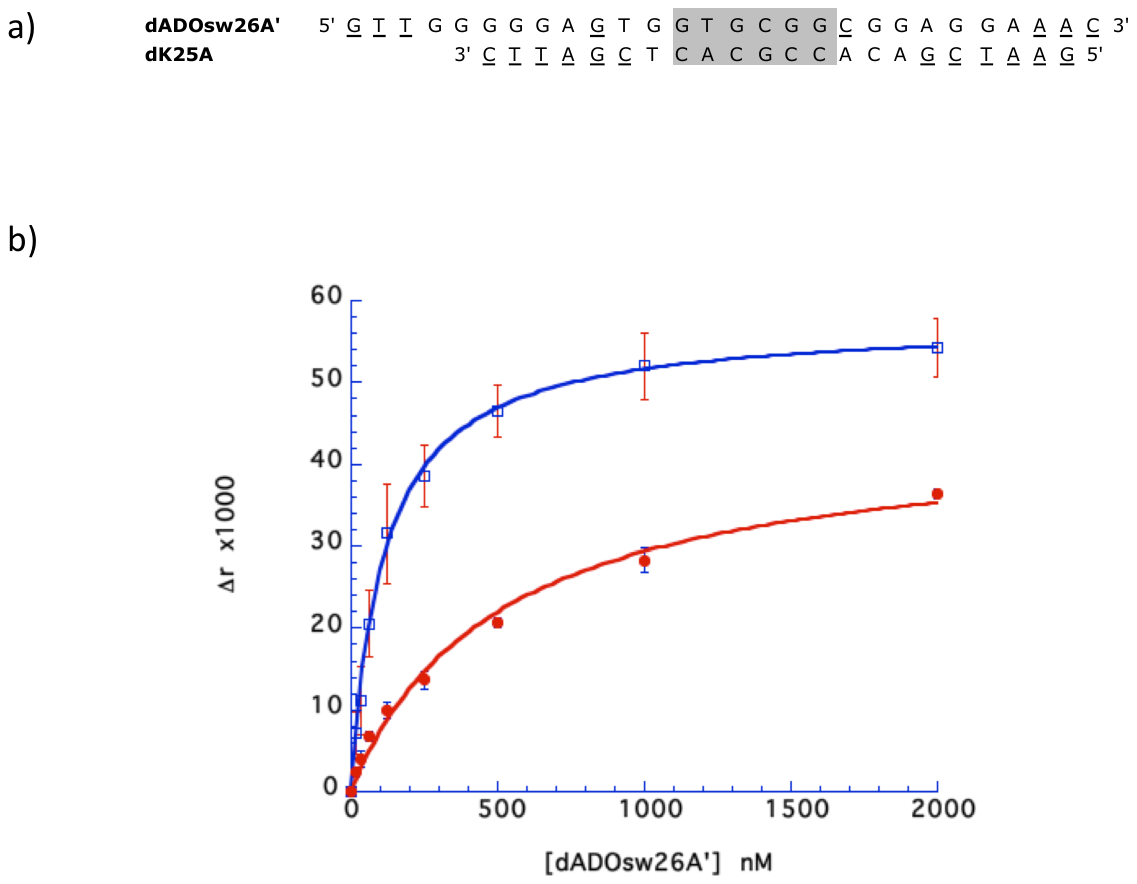

Fig 3) Determination of the affinity of the dADOsw26A’ adenoswitch for the Rox-dK25A aptakiss in the absence or presence of adenosine by fluorescence anisotropy measurement. 10 nM of the Rox-labeled-aptakiss-dK25A were incubated for 4 hours at 4°C with an increasing concentration of dADOsw26A’ ranging from 0 to 2 µM in the R buffer containing 10 mM of Mg^2+^(blue). Same experiment with increasing concentrations of dADOsw26A’ from 0 to 2 µM in presence of 1 or 2 mM adenosine (red).
